# Inhibitory properties of microalgal extracts on the *in vitro* replication of cyprinid herpesvirus 3

**DOI:** 10.1101/2021.03.05.434077

**Authors:** Fritzsche Stefanie, Blenk Patrik, Christian Jürgen, Castiglione Kathrin, Becker Anna Maria

## Abstract

Microalgae often stand out for their high biodiversity as well as their associated large number of potent bioactives. Therefore, they are interesting candidates as possible sources of antiviral substances, e.g. against cyprinid herpesvirus 3 (CyHV-3). Although this virus leads to high mortalities in aquacultures, there is no treatment available yet. Hence, ethanolic extracts produced with accelerated solvent extraction from six microalgal species *(Arthrospira platensis, Chlamydomonas reinhardtii, Chlorella kessleri, Haematococcus pluvialis, Nostoc punctiforme* and *Scenedesmus obliquus)* were examined in this study for inhibitory effects on viral replication. An inhibition of the *in vitro* replication of CyHV-3 in common carp brain cells could be confirmed for all six species, with the greatest effect for the *C. reinhardtii* and *H. pluvialis* extracts. At still non-cytotoxic concentrations viral DNA replication was reduced by over 3 orders of magnitude (> 99.9 %) each compared to the untreated replication controls, while the virus titers were at or even below the limit of detection. When pre-incubating cells and virus with *C. reinhardtii* and especially *H. pluvialis* extracts before inoculation, the reduction of viral DNA and virus titer was even stronger. Based on these results, an intervention in the initial replication steps like viral adsorption or membrane fusion is assumed. Moreover, a protection mechanism preventing the production of viral proteins and the assembly of mature virions is also possible. All in all, the results show that microalgae are a very promising source of natural antiviral substances against CyHV-3.

## Introduction

The breeding of common carp *(Cyprinus carpio)* as edible fish, with a worldwide commercial production of about 30 million tons, and koi as ornamental fish plays a key role in global aquaculture (Food and Agriculture Organization of the United Nations 2017). However, both – carp and koi – have been threatened worldwide by an animal disease caused by the cyprinid herpesvirus 3, CyHV-3, also called koi herpesvirus (KHV), belonging to the *Alloherpesviridae* family (Hedrick et al. 2000). CyHV-3 is an enveloped DNA virus with a double-stranded DNA in an icosahedral capsid of a total diameter of 170 to 230 nm (Ronen et al. 2003; Pokorova et al. 2005; Friedrich-Löffler-Institut 2016) that can cause the koi herpesvirus disease (KHVD) resulting in high mortality rates leading in turn to great financial damage (Gilad et al. 2003; Perelberg et al. 2003). Despite the wide spread of CyHV-3, no treatment against KHVD is currently available.

Commercial antivirals are based on the inhibition of virus adsorption and entry, protein synthesis and DNA replication or damage of mature virions. The majority of antivirals that are currently in use – such as acyclovir applied against herpes simplex viruses (HSV) – inhibit viral DNA synthesis (Modrow et al. 2010) by competing with natural guanosine triphosphates for chain incorporation and thus disturbing viral DNA replication (Villarreal 2001; Troszok et al. 2018). Key problems of these presently available antivirals are the substantial number of undesirable side effects, dose-dependent cell toxicity and the risk of developing resistance. Moreover, treatment of viral infections in animals faces problems due to possible drug residues, which preclude their use as food, and the issue of possible different cytotoxic effects of conventional human drugs on animals (Haetrakul et al. 2010; Modrow et al. 2010). Furthermore, additional factors such as antiviral application, its water solubility and stability have to be considered, in particular when treating fish in aquacultures. While current activities focus on the search for alternative antiviral substances, natural sources of antiviral agents are of growing interest for direct use as well as for further drug development for aquaculture (König 2007; Reichert et al. 2017).

Microalgae, with their enormous biodiversity, comprising an estimated number of species of up to several million as well as their ability to adapt to different habitats, are a potent source of bioactive compounds such as pigments, proteins, lipids and polysaccharides (Borowitzka 1995; Norton et al. 1996). For example, the already industrially used algal pigments astaxanthin, ß-carotene or phycobilins, are known for their range of bioactive effects, especially their antioxidative properties (Pulz and Gross 2004). Balzarini et al. (1991) discovered that plant (glyco)proteins have antiviral effects against human immunodeficiency virus (HIV) and human cytomegalovirus (HCMV) by preventing/inhibiting cell-virus fusion. Furthermore, microalgal sulfolipids, e.g. sulfoquinovosyldiacylglycerides (SQDG), are also regarded as potent antivirals (Naumann 2009). Gustafson et al. showed already in 1989 that SQDG extracted and purified from cyanobacteria impair the replication of HIV. In particular, sulphated polysaccharides of microalgal origin such as calcium spirulan and carrageenans show antiviral effects against HIV, influenza and especially against herpesviruses such as HSV (Witvrouw and De Clercq 1997; Rechter et al. 2006; Harden et al. 2009; Ahmadi et al. 2015). Furthermore, there is evidence that among the defense mechanisms that algae developed evolutionarily antiviral substances against *Phycodnaviridae*. Members of this virus family infect a variety of microalgae and have a similar morphology and replication cycle as herpesviruses (Chen and Suttle 1996; Thulke 2007). Thus, it is assumed that microalgal species contain defensive bioactives against viral infections that can also be effective against herpesviruses. Therefore, several studies have already reported on bioactives extracted from microalgal biomass that are effective against various viruses and especially against herpesviruses (Fabregas et al. 1999; Santoyo et al. 2012). However, only a few studies have been directed to antiviral substances against CyHV-3 in particular (Reichert et al. 2017; Haetrakul et al. 2018).

To examine microalgae for biologically active substances, the latter have to be extracted from the biomass to be tested for their activity. Accelerated solvent extraction (ASE) is often the method of choice when the target metabolites are at low concentrations and not of high stability. During ASE, increased pressures (3 to 200 bar), mostly elevated temperatures (up to 200 °C) and organic or aqueous solvents are used for the isolation of substances from solid samples (Luthria et al. 2004). The relatively low volume of required extraction solvent has the advantage of lower dilution of the extracted substances (Gey 2015) and additionally, sensitive metabolites are protected from degradation processes by the oxygen-free and low-light environment. Moreover, in comparison to conventional extraction methods, ASE not only requires shorter extraction times but also enables significantly higher extraction yields (Santoyo et al. 2009; Herrero et al. 2013). ASE has become a widely used and reliable technique for the extraction of various valuable products from natural sources, including phenolic compounds, polysaccharides, fats and pigments (Ligor et al. 2018; Nandasiri et al. 2019; Scott Chialvo et al. 2020). Also, the extraction of antiviral agents from microalgae using this technique has previously been reported (Santoyo et al. 2012).

In this study, ethanolic extracts from six microalgal species were prepared by ASE and their effect on the *in vitro* replication of CyHV-3 was investigated to identify novel antiviral substances against this pathogen which threatens one of the most important fish in aquaculture.

## Material and Methods

### Chemicals

Dulbecco’s modified Eagle medium (DMEM, Invitrogen Thermo Fisher Scientific Waltham, USA) was used as cell culture medium for cell maintenance in this work. Fetal calf serum (FCS) was obtained from Biochrom AG (Berlin, Germany). Analytical grade salts, titrants, ethanol (EtOH, ≥ 99.9 %), DMSO and HEPES (Pufferan, ≥ 99.5 %) were purchased from Carl Roth GmbH (Karlsruhe, Germany). PBS (pH 7.4) was prepared by mixing 137 mM NaCl with 2.7 mM KCl, 8.1 mM Na_2_HPO_4_ and 1.5 mM KH_2_PO_4_, all dissolved in ultrapure water (H_2_Ou) (Millipore, Darmstadt, Germany), autoclaved at 121 °C for 20 minutes and stored at room temperature (RT) in darkness until use. Antibiotic and antimycotic solution A5955 (with 10 000 units penicillin, 10 mg streptomycin and 25 μg amphotericin B per mL), Igepal (Nonident P40 Substitute) and Accutase^®^ were purchased from Sigma-Aldrich (St. Louis, USA). MTT (3-(4,5-dimethylthiazol-2-yl)-2,5-diphenyltetrazolium bromide) was purchased from SERVA Electrophoresis GmbH (Heidelberg, Germany).

### Cell cultures and virus stock

Common carp brain cells (CCB, passage 83, provided by the Friedrich-Loeffler-Institut (FLI), Greifswald, Germany) were maintained in 25-cm^2^ cell culture flasks (T-25, Sarstedt, Nümbrecht, Germany) in DMEM supplemented with 25 mM HEPES and 10 % FCS at 25 °C (Neukirch et al. 1999). CyHV-3 (KHV-TP 30) (Wang et al. 2015), isolated by Dr. Peiyu Lee (Taiwan, 2005), was provided by the FLI (Greifswald, Germany). CyHV-3 stocks were prepared by inoculation (multiplicity of infection, MOI = 0.2) of CCB cells seeded in DMEM with a density of 60 000 cells cm^-2^ in a cell disc (1000 cm^2^, Greiner Bio-One GmbH, Kremsmünster, Germany). Infected cells were incubated at 25 °C for three days post infection (p.i.) and replication was stopped by freezing the whole cell culture at ∡80 °C. Finally, frozen cells and medium were thawed, resuspended, aliquoted, introduced to titer determination and frozen at −80 °C for further use.

### Algal biomass

In this study, four different strains of chlorophyta *(Chlamydomonas reinhardtii, Chlorella kessleri, Haematococcus pluvialis, Scenedesmus obliquus)* and two cyanobacterial strains *(Arthrospira platensis, Nostoc punctiforme*) were used for extraction. The biomass was taken from an in-house microalgae biomass collection that had previously been produced in various research activities relating to microalgae (freeze-dried after harvesting and stored at −20 °C). For further information see Table S1 (Supporting Material).

### Preparation of ethanolic extracts

Biomass extracts were prepared using an accelerated solvent extractor (ASE 350, DIONEX, Sunnyvale, USA). The extraction was conducted using EtOH:H_2_O mixtures with different ratios at two different temperatures. For the basic extraction 95 % v/v EtOH and RT was chosen, whereas enhanced extracts were prepared using an increased water content (80:20 EtOH:H_2_O, v/v) and applying 100 °C. Accordingly, 500 mg of freeze-dried biomass was placed in extractions cells (V = 22 mL) and sealed with cellulose filters (Thermo Fisher Scientific, Waltham, USA) on both sides. Next, the extraction cells were filled with solvent and based on literature static extraction was performed for 30 min at a pressure of 100 bar for both extraction procedures (Santoyo et al. 2010; Herrero et al. 2013). N_2_ was used to purge the solvent from the cells into 60-mL vials that were closed with Teflon septa. To avoid contamination and carry-over of extracts the system was rinsed with solvent between extractions. Next, the extraction solvent was removed using a centrifugal evaporator (SPD131DDA, Thermo Fisher Scientific, Waltham, USA, RT, p = 10^-3^ bar). To minimize sample degradation due to oxidation, the dried extract residue was overlaid with N_2_ and stored in the dark at −20 °C until use. Each biomass extraction was performed in triplicates together with one blank sample (an empty extraction cell). The extraction yield was determined by dividing the exact dry weight of the extract residue by the initial dry biomass used for extraction. Afterwards, the dried extract residues were resuspended in non-toxic concentrations of DMSO (1 vol.-% at the lowest dilution level; continuously decreasing DMSO concentrations with increasing dilution) or EtOH (≤ 1 vol.-%) and selected volumes of cell culture medium (DMEM). Samples were additionally placed in an ultrasonic bath for 3 min (SONOREX, Bandelin, Berlin, Germany). In order to remove undissolved particles and to ensure sterility, redissolved extracts were filtered through syringe filters (0.22 μm, polyether sulfone, Sarstedt, Nümbrecht, Germany) in sterile 1.5-mL reaction vessels (Sarstedt, Nümbrecht, Germany) and kept at 8 °C in darkness until used.

### Evaluation of cytotoxic and antiviral effects of microalgal extracts

The cytotoxic potential of the extracts was examined using the colorimetric viability MTT assay. Briefly, CCB cells were seeded into 96-well plates with a density of 40 000 cells cm^-2^ in 100 μL DMEM supplemented with 25 mM HEPES, 0.5 M NaHCO_3_, 10 % FCS, 100 U mL^-1^ of penicillin G, 0.1 mg mL^-1^ of streptomycin sulphate and 0.25 μg mL^-1^ of amphotericin B and incubated at 25 °C for 24 h. Next, the medium was removed from the wells and replaced with 100 μL DMEM containing six consecutive dilutions (1:2) of the redissolved extracts (n = 6 for each concentration) and incubated for 72 h at 25 °C. Additionally, DMEM with 1 % DMSO/EtOH was used as 100 % viability controls (n = 6) and DMEM containing 20 % DMSO as 0 % viability controls (n = 6). After incubation, the medium was discarded and replaced with 100 μL of MTT solution, which was prepared from MTT (0.5 % w/v MTT in 0.9 % w/v NaCl solution) and DMEM in the ratio of 2:1 (v/v). The plate was incubated again for 4 h at 25 °C, followed by centrifugation at 3 220 g for 10 min. After discarding the supernatant, 20 μL of Igepal (0.4 %) was added to lyse the cells and the plate was incubated for 5 min under shaking at 1 000 rpm (Titramax 100, Heidolph, Schwabach, Germany). Next, formazan crystals were dissolved by adding 180 μL of DMSO and shaking the plate again for 5 min at 1 000 rpm. Finally, the absorbance of the samples and controls was measured at a wavelength of 570 nm using a plate reader (Multilabel Reader, Perkin Elmer, Waltham, USA). The average of the absorbance values of the 0 % viability control was subtracted from all other values which were then normalized to the growth control (100 % viability control). The determined cell viabilities were plotted against the corresponding extract concentrations and half-maximal effective concentrations (EC_50_) calculated using SigmaPlot and a *Four Parameter Logistic Curve*.

To evaluate the antiviral effects of the microalgal extracts CyHV-3 DNA was determined using a real-time qPCR (TaqMan) procedure as described by Gilad et al. (2004)with some modifications. Firstly, viral DNA was extracted from infected CCB cells using the DNeasy Blood & Tissue Kit (QIAGEN, Hilden, Germany) according to the manufacturer’s instructions. In addition, Proteinase K was used for an improved release of the nucleic acids and the breakdown of proteins during cell lysis. The obtained DNA extract (100 μL) was stored at −20 °C until further investigation and subjected to qPCR analysis using 96-well plates (Bio-Rad, Hercules, USA), the Thermocycler CFX connect system (Bio-Rad, Hercules, USA), and the temperature profile recommended by the manufacturer (see Table S2, Supporting Material). iTaq Universal Probes Supermix (Bio-Rad, Hercules, USA) was used with CyHV-3 specific primers and probes (Sigma-Aldrich, St. Louis, USA). Further information is given in the Supporting Material (Table S3). Glucokinase expression was used as an internal control and analyzed using glucokinase-specific primers and probes (Sigma-Aldrich, St. Louis, USA). A logarithmically diluted CyHV-3 standard series was prepared for the quantification of viral DNA. The half-maximal inhibitory concentration (IC_50_) of the extracts was determined based on the CyHV-3 DNA copy numbers, normalized to the values of the replication control and plotted against the extract concentrations by SigmaPlot (*Four Parameter Logistic Curve)*.

Infectious CyHV-3 viral particles were estimated using the endpoint dilution assay (50 % tissue culture infective dose assay, TCID_50_) as outlined by Reed & Muench (1938) and described previously by Mletzko et al. (2017) as well as Amtmann et al. (2020). The virus titer was registered as TCID_50_ mL^-1^.

To assess the influence of the extracts on the viral replication, the number of CyHV-3 DNA copies and the virus titers of the samples were compared with the values of the untreated virus replication control. To quantify this comparison, the logarithmic reduction factor R, which indicates the difference between two sample groups as orders of magnitude, was used. R was calculated according to equation 1.

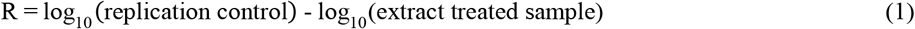

In order to assess whether the differences between replication control and samples are statistically significant, a one-way ANOVA (Excel) with a significance level of α = 0.05 was carried out.

Finally, to evaluate the strength of the antiviral effect compared to the cytotoxic effect, the selectivity index (SI) was determined using EC_50_ and IC_50_ according to equation 2.

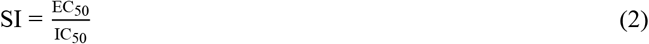

### Investigation of antiviral/inhibitory and cytotoxic properties of ethanolic extracts of various microalgae against CyHV-3 and CCB cells

Generally, the cytotoxic and inhibitory properties of the extracts were examined by incubation of CCB cells with or without inoculation with CyHV-3, after addition of consecutive (1:2) extract dilutions (n = 6) in 96-well plates. CCB cells were inoculated with a defined viral titer to obtain a MOI of 0.01.

For the investigation of antiviral properties of the microalgal extracts, CCB cells were seeded in 96-well plates (60 000 cells cm^-2^) and incubated at 25 °C for 24 h prior to the treatment. Extracts were tested in five to six different concentrations, obtained by consecutive dilutions (1:2) with DMEM as described above and mixed with virus stock to reach the appointed MOI. The concentration of the extracts was determined by taking the dry extract weight, the solvent volume and the dilution level into account and expressed in mg of dried extract per mL (mg_de_ mL^-1^). Firstly, the basic extracts (extracted at RT with 95 % v/v EtOH, n = 3) were added to the CCB cells together with the virus suspension. Samples were then incubated for 72 h at 25 °C. After incubation viral replication was stopped by freezing at −80 °C for at least 4 h and the samples were analyzed using qPCR (as described above) for CyHV-3 DNA copy numbers. In further experiments, extracts produced at an elevated extraction temperature (100 °C) using EtOH:H_2_O mixture (80:20, v/v) were examined. These enhanced extracts were diluted as described above and mixed with the virus stock to reach the same MOI. Extract/virus stock solutions prepared in this manner were added to the seeded CCB cells which then were incubated for 1 h at 25 °C. Since the period of 1 h is normally sufficient for viruses to penetrate the cells, the inoculum was discarded after this time. Cells were then overlaid with DMEM containing the same concentrations of the extracts and incubated for 72 h (25 °C) before viral replication was stopped by freezing as described above. The samples were analyzed using the same analytical procedures for CyHV-3 DNA copy numbers and viral titer. Additionally, MTT assays with the same concentrations of the extracts were performed to evaluate their cytotoxic effects on CCB cells. Samples containing the same virus load without extract addition were used as replication controls. Growth controls just with DMEM (without virus and extract) and extract controls containing the highest applied extract concentration but no virus in DMEM were also prepared in parallel. For each extract (n = 3) samples were analyzed using both qPCR and TCID_50_, resulting in a total of three biological replicates for viral DNA and titer per species.

### Influence of pre-treatment on the antiviral effects of microalgal extracts

To investigate the possible mode of antiviral action of the extracts, various set-ups of extract addition and inoculation of the cells were examined. The experiment was performed analogously to the procedure described above using CCB cells seeded with an identical density (60 000 cells cm^-2^) into 96-well plates and pre-incubated for 24 hours at 25 °C. Inoculation with CyHV-3 (1 h, 25 °C) was carried out using four different virus/extract addition procedures: (1) Cells were pre-treated with a previously defined, non-toxic concentration of extract in DMEM (1 h, 25 °C) and inoculated with CyHV-3 stock solution containing the same concentration of the extract. (2) The virus stock was mixed with the same non-toxic concentration of extract (1 h, 25 °C) and then used to infect CCB cells. In a third set-up, (3) the combination of the aforementioned cell and virus pre-treatment was tested and finally, for comparison, (4) cells were infected with a CyHV-3/extract mixture without any pre-incubation (neither cells nor extract). Regardless of the procedure, the supernatant used for inoculation was discarded after 1 h and the cells were overlaid with the corresponding extract/cell culture solution and incubated for 72 h at 25 °C before stopping the viral replication. Resulting samples were analyzed via qPCR and TCID_50_ for CyHV-3 DNA and titer.

## Results and discussion

### Extract preparation with the basic protocol

In this work, ASE was chosen for the extraction of potentially bioactive substances from microalgal biomass under defined conditions. Due to reports classifying EtOH as a safe and environmentally friendly solvent suitable for the extraction of natural value products from microalgae (Gallego et al. 2018; Molino et al. 2018), an EtOH:H_2_O mixture with 5 % v/v water content was chosen. Considering a possible thermolability of target molecules, the extraction process was initially performed at RT (*basic extraction*). As summarized in Table 1, the obtained yields varied between 2.8 % and 22 % depending on the biomass. There was no clear correlation between extraction yield and taxonomic affiliation of the organisms, since both the best and the worst extraction were achieved with species from the division of chlorophyta.

**Table 1:** Yield (% w/w, based on dry weight) of biomass extract obtained by accelerated solvent extraction using different temperatures and solvent compositions

| Species | Taxonomy | Extraction method | Yield, % w/w |
| --- | --- | --- | --- |
| *A. platensis* | Cyanobacteria | basic | 5.6 ± 0.2 |
|  |  | enhanced | 32 ± 0.8 |
| *C. kessleri* | Chlorophyta | basic | 20 ± 0.8 |
|  |  | enhanced | n.e. |
| *C. reinhardtii* | Chlorophyta | basic | 18 ± 0.6 |
|  |  | enhanced | 29 ± 0.8 |
| *H. pluvialis* | Chlorophyta | basic | 22 ± 0.3 |
|  |  | enhanced | 29 ± 1.8 |
| *N. punctiforme* | Cyanobacteria | basic | 8.5 ± 0.2 |
|  |  | enhanced | 11 ± 0.4 |
| *S. obliquus* | Chlorophyta | basic | 2.8 ± 0.5 |
|  |  | enhanced | n.e. |

Basic extraction: RT, 95:5 EtOH:H<sub>2</sub>O, v/v
Enhanced extraction: 100 °C, 80:20 EtOH:H<sub>2</sub>O, v/v
n.e.: not examined

### Effects of the basic extracts on virus replication and cellular vitality

To obtain a first assessment of possible inhibitory effects of the basic extracts on viral replication, CCB cells were infected with CyHV-3 along with different extract concentrations and were incubated for 72 h at 25 °C without removing the viral supernatant after inoculation. In this set-up, the samples were only analyzed by characterizing the CyHV-3 DNA copy number, always in comparison to the replication control (without extract addition). With this fast-forward test, viral replication could be monitored, but not the quantity of infectious viral particles. Therefore, samples showing a possible inhibiting effect on viral DNA replication were additionally analyzed regarding their viral titer using the TCID_50_ method. To exclude that a reduced viral replication was merely a result of damaged host cells, the cytotoxic influence of the extract on CCB cells was examined using the same extract concentrations.

Since the yields of the basic extracts differed by up to a factor of 9, the maximum concentrations that could be used for the first effectiveness evaluation were very different, as shown in Figure 1. Furthermore, in case of *H. pluvialis* and *A. platensis,* an additional dilution factor has to be considered because of the maximal applicable solvent concentration and an increased solvent use due to poor solubility of these basic extracts. We therefore examined whether the extract concentrations were sufficient to determine IC_50_, EC_50_ and SI, since these values allow for an evaluation of the strength of the antiviral effect in comparison to the cytotoxic effect. In general, the highest possible SI should be aimed for. Acyclovir, which is often used against herpes viruses, for example, can achieve an SI well over 100 for HSV-1 (Lucero et al. 2006; Defer et al. 2009). The IC_50_, and EC_50_ values were determined on the basis of the corresponding dose-response curves (Figure S1, Supporting Material) and are summarized in Table 2. Samples treated with the basic extracts of *A. platensis* and *H. pluvialis* showed no or only minor differences in viral DNA concentration compared to the replication control (Figure 1 a,b). The maximum concentrations of 0.6 mg_de_ mL^-1^ and 2.2 mg_de_ mL^-1^, respectively, were not sufficient to reach clear effects on the cell vitality or viral replication. Therefore, neither EC_50_ nor IC_50_ could be determined. In contrast, using *C. kessleri, N. punctiforme*, *S. obliquus* and *C. reinhardtii* extracts both cell viability and viral replication of CyHV-3 were affected in a dose-dependent manner (Figure 1 c-f, S1). Depending on the biomass used, these effects showed up differently. Due to the strongest cytotoxic effect, only a low SI could be determined for the extract of *S. obliquus* (Table 2). The calculated SI for *C. kessleri* was also comparatively low, since both inhibitory and cytotoxic effects occurred only at high concentrations resulting in high IC_50_ and EC_50_ values. The extracts of *C. reinhardtii* and *N. punctiforme* showed a comparable antiviral activity to the extract from *S. obliquus,* but at the same time an approximately 3-fold lower cytotoxicity, which resulted in higher SI values of 4.3 and 4.1, respectively (Table 2). A more detailed analysis of the antiviral activity at extract concentrations with only moderate effects on the cell viability (Table 3) also showed that the extracts from *C. reinhardtii* and *N. punctiforme* were the most promising ones, which led to a reduction of viral DNA by 0.7 and 1.6 orders of magnitude, respectively. However, in comparison to *N. punctiforme,* the viral titer was reduced stronger (2.2 orders of magnitude in comparison to 1.4) when using the extract of *C. reinhardtii.* Notably, also the other basic extracts that showed antiviral activity led to a reduction of both the viral DNA copy numbers and the viral titers under these conditions.

**Figure 1:**
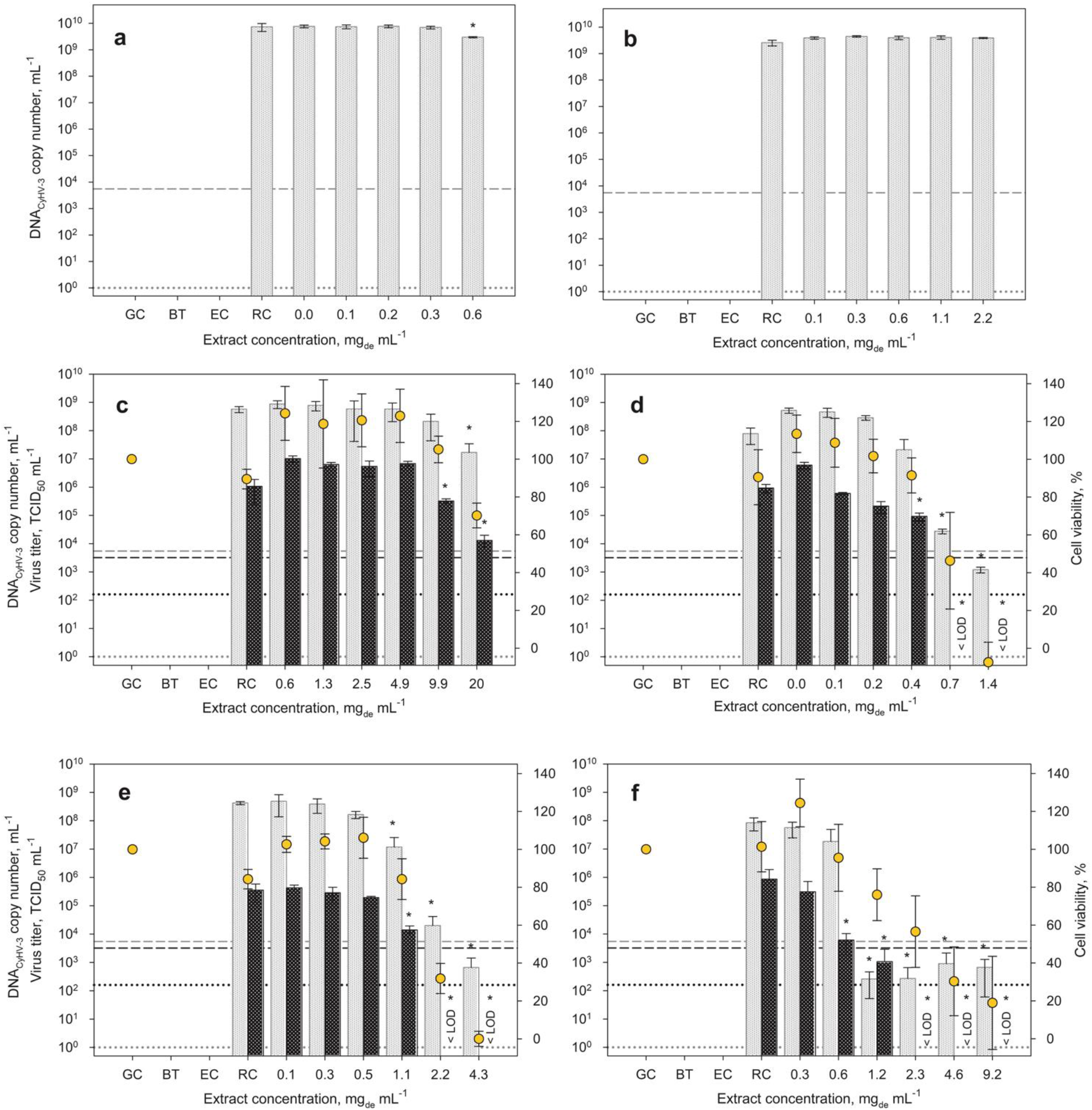
Effect of various concentrations of ethanolic extracts of *A. platensis* (a), *H. pluvialis* (b), *C. kessleri* (c), *S. obliquus* (d), *N. punctiforme* (e) and *C. reinhardtii* (f) on viral replication of CyHV-3. Each bar of viral DNA copy numbers (shown in gray) and infectious viral titer (shown in black) represents the mean of three biological replicates ± SD. In addition, the vitality of CCB cells normalized to the growth control is shown. Initial viral DNA copy number and virus titer are shown by dashed lines, while the respective limit of quantification is represented by dotted lines. RC – represents replication controls with the CCB cells infected with the same virus load without extract addition; GC – corresponds to growth controls (mock infected without virus and extract) and EC – displays extract controls containing the highest applied extract concentration but no virus in cell culture medium. If the virus titers could no longer be detected using TCID_50_, this is marked with < LOD (limit of detection). *p ≤ 0.05 (compared to replication control, RC).

**Table 2:** Cytotoxicity (EC_50_) and antiviral activity (IC_50_) of microalgal extracts obtained with the basic extraction protocol. For associated dose-response curves see Figure S1 (Supporting Material).

| Species | EC<sub>50</sub>, mg<sub>de</sub> mL<sup>-1</sup> | IC<sub>50</sub>, mg<sub>de</sub> mL<sup>-1</sup> | SI, - |
| --- | --- | --- | --- |
| *A. platensis* | - | - | - |
| *C. reinhardtii* | 1.7 ± 0.7 | 0.4 ± 0.1 | 4.3 ± 2.0 |
| *C. kessleri* | 24 ± 5.2 | 8.4 ± 6.0 | 2.9 ± 2.7 |
| *H. pluvialis* | - | - | - |
| *N. punctiforme* | 1.8 ± 0.1 | 0.4 ± 0.1 | 4.1 ± 0.5 |
| *S. obliquus* | 0.6 ± 0.1 | 0.4 ± 0.1 | 1.4 ± 0.7 |

EC<sub>50</sub>: 50 % effective concentration, IC<sub>50</sub>: 50 % inhibitory concentration, SI: selectivity index

**Table 3:** Effects of *C. kessleri, N. punctiforme, S. obliquus* and *C. reinhardtii* extracts obtained with the basic extraction protocol on CyHV-3 replication. Reduction of viral DNA copy number and virus titer was analyzed at concentrations at which cell viability was only slightly affected.

| Species | Concentration, mg<sub>de</sub> mL<sup>-1</sup> | Cell viability, % | Reduction of DNA, orders of magnitude (%) | Reduction of virus titer, orders of magnitude (%) |
| --- | --- | --- | --- | --- |
| *C. kessleri* | 9.9 | 105 | 0.4 (60) | 0.5 (68) |
| *N. punctiforme* | 1.1 | 86 | 1.6 (98) | 1.4 (96) |
| *S. obliquus* | 0.4 | 90 | 0.6 (75) | 1.0 (90) |
| *C. reinhardtii* | 0.6 | 96 | 0.7 (80) | 2.2 (> 99) |

Due to the significant reductions of viral DNA copies, a negative influence on the replication of CyHV-3 by the extracts of *C. kessleri*, *S. obliquus, N. punctiforme* and *C. reinhardtii* can be assumed, despite some cytotoxic effects of the latter (Table 2). These results are in good agreement with the already reported antiviral effects of *N. punctiforme* and *S. obliquus* extracts against enveloped DNA viruses (Kanekiyo et al. 2005; Naumann 2009; Singab et al. 2018). As described in these studies, it is possible that polysaccharides (e.g. nostoflan), short-chain free fatty acids and especially glycolipids such as SQDG play a role in the inhibition of virus-cell adsorption and membrane fusion and thus infection. It is believed that similar compounds also mediate the antiviral effects of extracts from *C. kessleri* (Plaza et al. 2012; Yao et al. 2015). In this work, virus titer reductions below the limit of detection were observed with extracts from *S. obliquus*, *N. punctiforme* and *C. reinhardtii* (Figure 1 d-f). It is possible that amphiphilic molecules, such as monoglycerides, contained in aqueous-ethanolic extracts damaged the lipid envelope of virions and therefore even inactivated the viruses as it was shown for *S. obliquus* extracts in previous studies (Thromar et al. 1994; Ishaq et al. 2016).

### Extract preparation with an enhanced protocol

As already mentioned, the extraction conditions play a major role in the extraction efficiency and the substances extracted, determined by the biomass used. Depending on the selected solvent and the extraction temperature, different substances can be dissolved from the biomass to different extent. We therefore started a second extraction run to test how modified extraction conditions might affect and improve the yield and the virus inhibiting effect of the extracts. In this second set-up, a focus was put on *C. reinhardtii* and *N. punctiforme* after the most promising effects were observed for the corresponding basic extracts. The extraction of *A. platensis* and *H. pluvialis* was also re-examined to test whether other extraction conditions would result in higher applicable concentrations and perhaps other bioactive effects, as these two microalgae has shown antiviral properties against various viruses before (Santoyo et al. 2012; Nuhu 2013). Especially *A. platensis* is known to be a source of extra- and intracellular polysaccharides that were identified in several studies as potent antiviral substances against e.g. HSV and also CyHV-3 (Hayashi et al. 1993; Reichert et al. 2017; Singab et al. 2018). Thus, it was considered that the parameters chosen for the basic extraction might have not been suitable to extract these metabolites. Therefore, the water content of the extraction solvent was increased from 5 to 20 % (v/v) to facilitate the extraction of more polar compounds. Additionally, the influence of an elevated extraction temperature (100 °C) for the isolation of anti-CyHV-3 bioactives was evaluated in this second experimental set-up called *enhanced extraction*.

For all four species, the extraction yield could be raised by factors between 1.3 and 5.7 by applying the enhanced protocol as can be inferred from Table 1. These results suggest that indeed more metabolites were extracted from the applied biomass when using an increased water content in the solvent and a higher extraction temperature, which is in good agreement with previously published data (Santoyo et al. 2010; Tolivia et al. 2013). Moreover, the improved extraction procedure enabled us to perform further studies on the antiviral properties of the selected species with higher concentrations of the extracts.

### Effects of the enhanced extracts on virus replication and cellular vitality

Applying the enhanced extracts CyHV-3 infection experiments were performed again using the above-explained procedures, but now with a deviation affecting the incubation post infection. It cannot be ruled out that virus particles, whose penetration has been initially prevented by the extract, subsequently infect cells anyway after prolonged exposure to the cells because of a possible weakening extract effect due to for example oxidizing impacts. In such a case, higher infectious titers would be measured than induced by a fresh extract. Therefore, based on previous studies (Hayashi et al. 1993; Hernández-Corona et al. 2002a; Santoyo et al. 2012) and since we assumed that 1 h is sufficient for the virus to enter the cells, further experiments were carried out without constant virus exposure. Thus, to remove unabsorbed viruses after the initial infection phase the infectious supernatant of all samples and also the supernatant of the replication controls was discarded after inoculation. Subsequently, fresh cell culture medium with the appropriate extract concentrations was added to the cells. As a result, using the enhanced extracts, a dose-dependent inhibition of viral replication could now be observed for the extracts of all four species (Figure 2). Although, no effects on the CyHV-3 replication were seen in the previous experiments for *A. platensis* and *H. pluvialis*, now inhibition was registered when using the enhanced extraction. Applying the *A. platensis* extract at high concentrations, a reduction in CyHV-3 DNA copy numbers occurred up to approximately 5 orders of magnitude (Figure 2 a). Hence, to verify the data, the infectious viral particles were analyzed using TCID_50_ besides the characterization of viral DNA. For higher extract concentrations viral titers lay even below the limit of detection of the assay, corresponding to a reduction of over 5 orders of magnitude. However, dosedependent increasing cytotoxic effects of the *A. platensis* enhanced extract on the CCB cells were observed. Thus, the EC_50_ and IC_50_ values calculated from these results led to a relatively low SI (Table 4). As described above for *A. platensis,* also for *H. pluvialis* higher extract concentrations could be applied for the antiviral studies due to the improved extraction procedure. Consequently, inhibiting effects were also revealed for this alga. In this case, significant reductions of viral DNA of up to 4 orders of magnitude (99.99 %) occurred, while the viral titer was no longer detectable (Figure 2 d). It was noticeable that no significant signs of damage to the CCB cells were detectable via the MTT assay even at such elevated extract concentrations. Up to concentrations of 7.4 mg_de_ mL^-1^ the viability of CCB cells was actually exceeding the growth control and decreased to a minimum of yet 90 % at the highest extract concentration used here. These high viabilities at elevated extract concentration led to an increased EC_50_ value, resulting in a comparably high SI of 60 (Table 4). Overall, using enhanced *A. platensis* and *H. pluvialis* extracts and the improved procedure, viral replication was significantly inhibited also without cytotoxic effects (Table 5). For both algae the dose-dependent inhibitory effect on CyHV-3 replication is in good agreement with the above-mentioned data for polysaccharides from *A. platensis* and herpesviruses such as HSV_1, human cytomegalovirus (HCMV) and human herpesvirus 6 (HHV6), where the interference with viral adsorption and entry into the target cells was suggested as a mode of action (Rechter et al. 2006; Singab et al. 2018). Also, for extracts of *H. pluvialis* an antiviral effect against HSV-1, where polysaccharides and short-chain free fatty acids were suggested as possible antiviral substances, was reported (Santoyo et al. 2012). Since, due to the chosen extraction conditions, polar lipids and polysaccharides could also be extracted in this work, a similar active agent and mechanism of action can be assumed. Furthermore, the data presented for these two species showed in which manner the selection of the experimental conditions and analysis methods could affect the outcome of such screening studies. If only the preliminary experiments with *A. platensis* and *H. pluvialis* were considered, the inhibitory activity of their extracts on CyHV-3 would remain unrecognized, due to the lower tested extract concentrations and the lack of effects on the viral DNA. However, when varying the experimental conditions and extending the analysis to the viral titer, very promising results regarding an antiviral effect against CyHV-3 could be registered.

**Figure 2:**
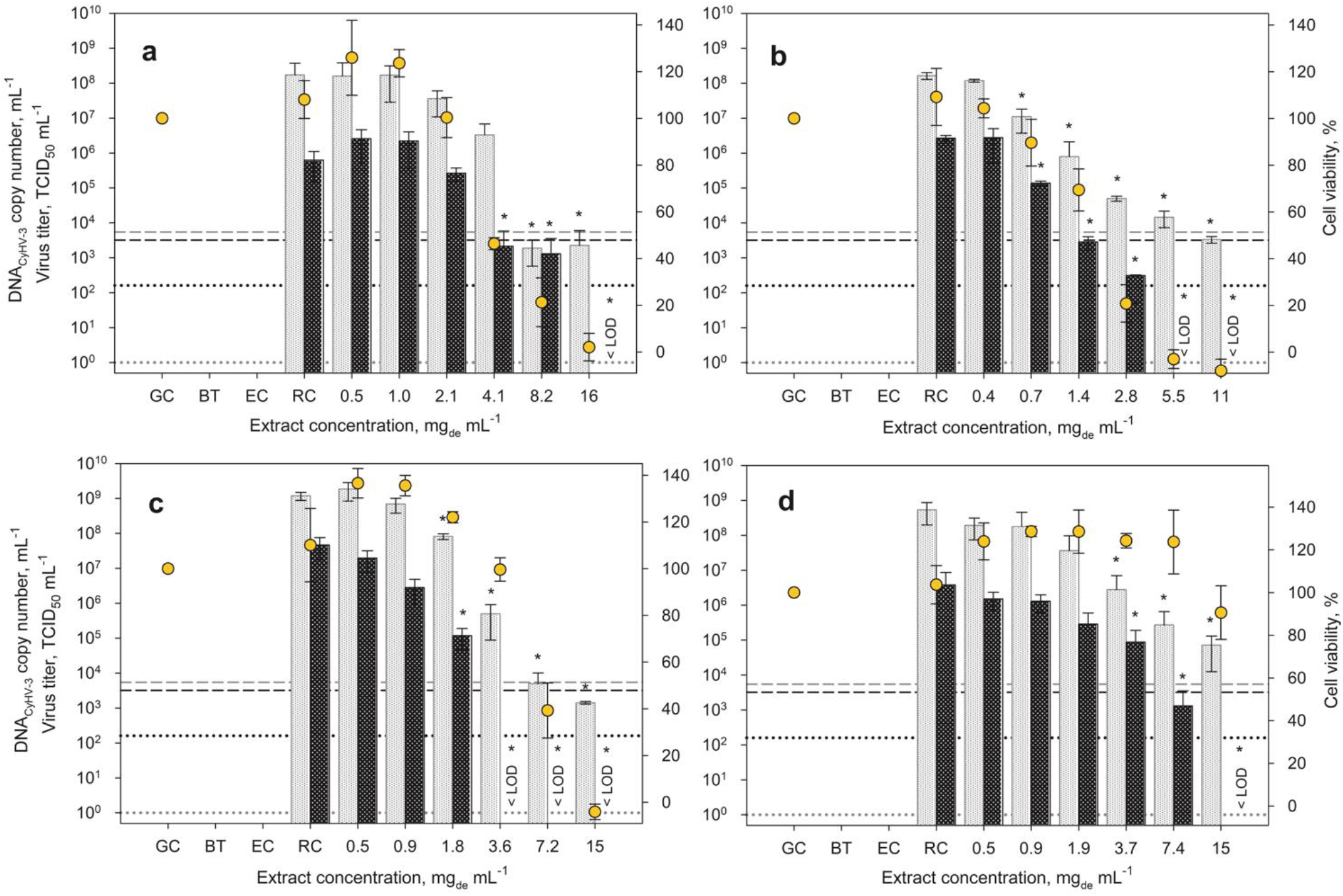
Effect of ethanolic extracts of *A. platensis* (a), *N. punctiforme* (b), *C. reinhardtii* (c) and *H. pluvialis* (d) on viral replication of CyHV-3. CCB cells were infected with CyHV-3, treated with extract during and after inoculation and were incubated at 25 °C for 1 h. Infectious supernatant was replaced with extract (in medium) and cells were incubated at 25 °C for 72 h. Each bar of viral DNA copy numbers (shown in gray) and infectious viral titer (shown in black) represents the mean of three biological replicates ± SD. In addition, the vitality of CCB cells normalized to the growth control is shown. Initial viral DNA copy number and virus titer are shown by dashed lines, while the respective limit of quantification is represented by dotted lines. RC – represents replication controls with the CCB cells infected with the same virus load without extract addition; GC – corresponds to growth controls (mock infected without virus and extract) and EC – displays extract controls containing the highest applied extract concentration but no virus in cell culture medium. If the virus titers could no longer be detected using TCID_50_, this is marked with < LOD (limit of detection). *p ≤ 0.05 (compared to replication control, RC).

**Table 4:**
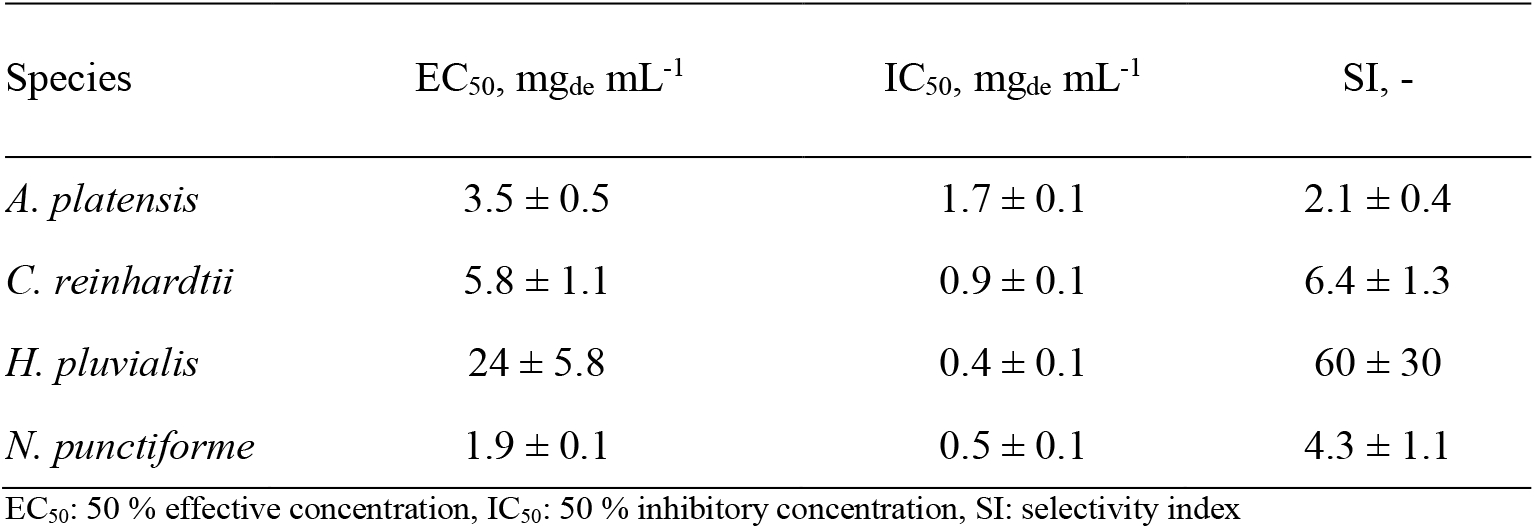
Cytotoxicity (EC_50_) and antiviral activity (IC_50_) of microalgal extracts obtained with the enhanced extraction protocol. For associated dose-response curves see Figure S2, Supporting Material.

**Table 5:** Effects of *A. platensis, C. reinhardtii, H. pluvialis,* and *N. punctiforme* extracts obtained with the enhanced protocol on CyHV-3 replication. Reduction of viral DNA copy number and virus titer was analyzed at concentrations at which cell viability was not affected.

| Species | Concentration, mg<sub>de</sub> mL<sup>-1</sup> | Cell viability, % | Reduction of DNA, orders of magnitude (%) | Reduction of virus titer, orders of magnitude (%) |
| --- | --- | --- | --- | --- |
| *A. platensis* | 2.1 | 100 | 0.7 (63) | 0.4 (58) |
| *C. reinhardtii* | 3.6 | 100 | 3.4 (> 99.9) | > 4 (> 99.99) |
| *H. pluvialis* | 7.4 | > 100 | 3.3 (> 99.9) | 3.5 (> 99.9) |
| *N. punctiforme* | 0.7 | 90 | 1.2 (94) | 1.3 (95) |

Furthermore, the influence of the changed extraction conditions was examined for *N. punctiforme* and *C. reinhardtii*. As with the basic extracts, a dose-dependent reduction of replicated viral DNA could be detected for both enhanced microalgae extracts (Figure 2 b,c). Moreover, a steadily decreasing titer of infectious viral particles was recognizable with increasing extract concentrations. Even at concentrations where replication of viral DNA was not yet affected, a reduced titer was found. It is noticeable that the IC_50_ values of the enhanced extracts could not be improved for both organisms. Quite the contrary, in the case of *C. reinhardtii* it was even slightly worse (Tables 2 and 4). Thus, the enhanced extraction led to higher concentrations and biomass yields, but to a lower inhibiting effect. This underpins the fact that with crude extracts an increased extraction does not necessarily correspond to a stronger effect. Nevertheless, whereas the SI values of the basic and enhanced extracts from *N. punctiforme* were comparable, the SI of the enhanced *C. reinhardtii* extract was improved by a factor of 1.5 since a reduced cytotoxic effect overcompensated the slight loss of antiviral activity. Table 5 shows that, among all considered microalgae in this study, the greatest inhibitory effect without damaging the CCB cells could be reached with the *C. reinhardtii* extract resulting in a reduction of viral DNA by 3.4 (> 99.9 %) and viral titer by over 4 (> 99.99 %) orders of magnitude. However, while no analysis of the extract composition of *C. reinhardtii* was performed in this work, there is some data published in peer-reviewed literature regarding bioactive compounds contained in extracts of this species. For example, Choudhary et al. (2018) determined that ethanolic aqueous extracts of *C. reinhardtii* consisted to almost 70 % of carbohydrates with a high sulphate content, based on which the existence of sulphated polysaccharides in this green alga could be proven afterwards (Choudhary et al. 2018; Kamble et al. 2018; Vishwakarma et al. 2019). In addition, Yao et al. (2015) showed a high content of glycolipids for isopropanolic *C. reinhardtii* extracts. Mattos et al. (2011) assumed the participation of algal glycolipids in antiviral activity against HSV-1 based on the prevention of the initial virus adsorption and membrane fusion with the cells. Therefore, it is possible that the inhibitory effects of the ethanolic *C. reinhardtii* extract against the replication of CyHV-3 observed here also result from (sulphated) polysaccharides and glycolipids preventing the initial steps of viral replication. Although various bioactive effects have been described in literature for extracts of *C. reinhardtii*, no inhibitory impact on the replication of viruses have been reported so far apart from the results obtained no infectious viral particles could be detected in this work. Again, this suggests that in studies regarding possible bioactive metabolites, both the choice of experimental conditions and the methods of analysis play an important role. Therefore, it is possible that *C. reinhardtii* was not yet considered as an interesting candidate as a source of antiviral compounds because, for example, other extraction methods or only one-sided analyses were used for evaluation and the corresponding data was not published. Hence, when finding no antiviral effects of extracts of specific species on selected viruses instantly these species should not be excluded from further investigation prematurely.

From the point of view of a future application and further studies, especially non-cytotoxic extract concentrations that are showing inhibitory effects on the viral replication are of great interest. With this in mind, the *C. reinhardtii* extract inhibited viral replication the most with unaffected cell viability (Table 5). On the other hand, the enhanced extract of *H. pluvialis* showed the lowest cytotoxicity and a high inhibitory influence on CyHV-3 which led to a relatively high SI of 60, which is significantly higher than the SI of 6.4 reached with *C. reinhardtii* (Table 4). However, the latter might be optimized in further research for example by the isolation and purification of the biologically active compounds. In general, biological extracts contain a variety of compounds, meaning that it is always unclear which biological effects are caused by which molecules. Thus, it has always to be analyzed whether a measured cytotoxic activity is a feature of the same molecule as the antiviral activity in order to avoid excluding active compounds from screenings only due to high cytotoxic effects of their crude extracts (Wildman 2003). Therefore, the isolation of the active antiviral or inhibitory agent or rather substance class might help to eliminate substances which affect the host cells cytotoxically and therefore improve the SI accordingly.

Since the SI of antiviral extracts and substances depends not only on the species, extraction method and the degree of purification, but also on the virus and the host cells used, cross-study comparisons must be done with caution and are better suited to comparing orders of magnitudes than concrete values. In a similar study performed with hot water extracts from *A. platensis,* the replication of HSV-1 and HSV-2 was inhibited by 50 % at a concentration of 0.3 mg mL^-1^ showing IC_50_ values in the same order of magnitude and an SI of 26 (Hernández-Corona et al. 2002b; Singab et al. 2018). However, when using purified products such as sulphated polysaccharides and SQDG, significantly lower IC_50_ values were achieved where replication of HSV and HCMV was inhibited by 50 % at concentrations in the range of μg mL^-1^ (Naumann 2009; Ahmadi et al. 2015). Exopolysaccharides extracted and purified from *A. platensis* resulted in an IC_50_ of less than 2.1 μg mL^-1^ but also in a low SI of 3 for CyHV-3 (Reichert 2016). Using *H. pluvialis* extracts, 99 μg mL^-1^ of the polysaccharide-rich fraction reduced virus infectivity by 50 % resulting in an SI of 19 (Santoyo et al. 2012).

### Influence of pre-treatment on the antiviral activity of the enhanced extracts

In order to obtain information about the virus replication phase affected by the inhibiting extracts and thus confirming the presumed mechanism of action of the positively tested extracts, different pre-incubation set-ups were examined. Either CCB cells or CyHV-3 suspensions were pre-incubated for 1 h with the selected concentrations of extracts before inoculation. In addition, a combined mode was examined, where both, cells and virus suspension, were pre-incubated with the extracts. After this pre-incubation, the assays for antiviral activity were carried out as described above. Using this strategy, it should be possible to differentiate, whether the inhibiting agents influence either the cells or the viral particles or even both. For comparison, samples with no extract pre-incubation (corresponding to the previous set-up for the investigation of antiviral properties of the extracts) and the untreated replication controls were prepared along with the other samples.

The results of the pre-treatment experiment with the enhanced extracts from *A. platensis*, *N. punctiforme, C. reinhardtii* and *H. pluvialis* are illustrated in Figures 3 a-d. For all samples where one of the four extracts was applied, varying inhibitory effects on viral DNA and titer were observed. Depending on whether cells, the virus suspension, both or neither of them were pre-incubated with the corresponding extract, the reductions in viral DNA or titer occurred to different degrees. Thus, while the least effect on viral infection was always observed when no pre-incubation was applied, the pre-treatment of both cells and virus prior to the inoculation with respective extracts resulted in significantly lower viral DNA and titer for all four species. When pre-incubating the extracts of *A. platensis* and *N. punctiforme* with either cells or virus suspension, only a slight influence on the number of copies of viral DNA was registered. Simultaneously, the same use of these extracts caused a more pronounced reduction of infectious virus particles compared to viral DNA – as it has also been observed in previous experiments. The application of *C. reinhardtii* extract to either cells, virus or both led to a reduction of viral DNA and viral titer below the limit of detection. However, even stronger effects were observed for each scenario in this set-up when using the extract of *H. pluvialis.* All in all, a more pronounced interference with viral replication could be recognized when the cells were in contact with the extract before inoculation. Consequently, it can be assumed that the extract affects components of the cells like impairing viral adsorption to the cells. Therefore, it is possible that the inhibition of the virus entry by sulphated polysaccharides as reported by Baba et al. (1988) and Hayashi et al. (1996) also contributes to the effects observed in this study. However, the data obtained especially for the combined pre-incubation set-up suggest that the antiviral effect is not solely based on the inhibition of virus binding to the host cells, as viral replication was much more impeded in the combined mode in comparison to just treating the cells. Most likely, also following replication steps such as virus assembly or release can be affected by the extracts used in this work. Furthermore, since infectious virus particles were also not detected after the virus stock was pre-treated with the extracts of *C. reinhardtii* and *H. pluvialis*, some virucidal properties of the latter can be assumed for these two extracts. Thus, the inhibitory effects on the replication seem to be a combination of impeded entry, possibly triggered cell defense mechanisms affecting the virion production and some virucidal properties. Nevertheless, a clear designation of mode of action of these extracts is not yet possible based only on the data presented here. Altogether, the pre-incubation data confirmed the inhibitory effects of the enhanced extracts of *A. platensis, N. punctiforme*, *C. reinhardtii* and *H. pluvialis* and form a basis for identifying the bioactive substances and their mechanisms of action in further studies.

**Figure 3:**
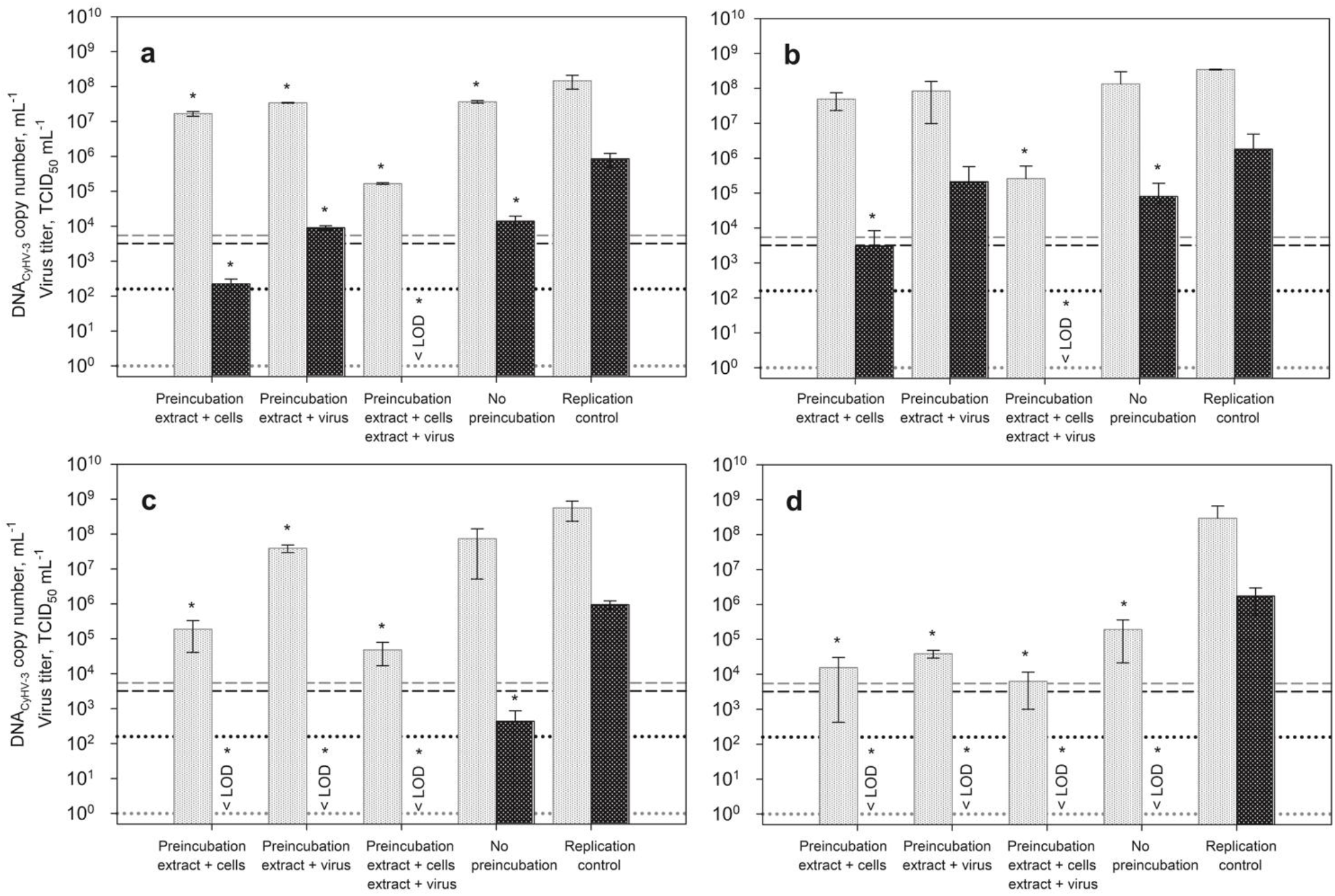
Effect of different pre-incubation modes with *A. platensis* (a), *N. punctiforme* (b), *C. reinhardtii* (c) *and H. pluvialis* (d) extract (concentrations: 2.1 mg_de_ mL^-1^, 0.7 mg_de_ mL^-1^, 1.8 mg_de_ mL^-1^ and 7.1 mg_de_ mL^-1^) on viral replication of CyHV-3. Either cells, virus suspension or both were pre-incubated with the ethanolic extracts before inoculation with CyHV-3 at 25 °C for 1 h. Infectious supernatant was replaced with extract (in medium) and cells were incubated at 25 °C for 72 h. The viability of the cells was approximately 100 % for each experiment, only for the incubation with *H. pluvialis* the cell viability was slightly affected and reduced to 90 %. Each bar of viral DNA copy numbers (shown in gray) and infectious viral titer (shown in black) represents the mean of three biological replicates ± SD. In addition, the vitality of CCB cells normalized to the growth control is shown. Initial viral DNA copy number and virus titer are shown by dashed lines, while the respective limit of quantification is represented by dotted lines. If the virus titers could no longer be detected using TCID_50_, this is marked with < LOD (limit of detection). *p ≤ 0.05 (compared to replication control, RC).

## Conclusion

In summary, ethanolic extracts of six different chlorophyta and cyanobacteria obtained by ASE inhibited the *in vitro* replication of CyHV-3, suggesting bioactives with antiviral properties against the selected virus present in these extracts. Therefore, the work presented here confirms the immense potential of the phototrophic microorganisms when searching for such agents. In general, antiviral effects of all extracts were observed after their application in the range of lower mg_de_ mL^-1^. We could demonstrate that ASE enables the extraction of bioactive substances and that the application of elevated temperature for the isolation process did not negatively affect the antiviral properties of the extracts tested here. On the contrary, elevated temperature and water content in the extraction solvent led to a higher extraction yield and enabled the identification of antiviral effects even for extracts of the species which, based on the preliminary experiments, seemed not to have such properties. Using the enhanced extracts of the four more intensively studied species – *A. platensis*, *N. punctiforme*, *C. reinhardtii* and *H. pluvialis* – a substantial reduction in virus replication (based on both viral DNA and titer) could be proven for all of them even for high cell viabilities. Within the investigated species, the clearest antiviral effects were registered for the *C. reinhardtii* and H*. pluvialis* extracts with the highest SI values calculated in this study. The data collected in this work suggest that this inhibitory effect is based on the interference of bioactive substances with the attachment of virus particles to the target cells, preventing viral entry as well as a possible activation of some cell defense mechanisms impeding the production of viral proteins and thus in a consequence the assembly of mature virions. To the best of our knowledge, this is the first report of antiviral properties of extracts of *C. reinhardtii.* Based on the experiments performed in this work, as well as on the data published previously on sulphated polysaccharides, it can be assumed that these substances might be responsible for the effects observed here on the replication of CyHV-3. However, according to literature also other compounds such as free fatty acids or polar (glyco)lipids could be involved. Furthermore, our work shows that the quantification of the actual infectious virus titer in addition to the viral DNA copy number is of great importance in order to assess the antiviral properties of such extracts. While in some cases viral DNA could still be produced by the extract treated and CyHV-3 infected cells, no infectious viral particles could be detected via titration assay. Moreover, a titer reduction under extract application was often more pronounced in comparison to the reduction of viral DNA. Thus, if only viral DNA was monitored the antiviral properties of some extracts might have remained unnoticed. Since a drop in the titer below the initially added values was observed for most extracts can also have virucidal effects, at least for the virus investigated here. Nevertheless, further research and the identification of the antiviral substances and their modes of action is necessary for the phototrophic species characterized in this study. In particular, further investigation of the more closely studied microalgal species *A. platensis, H. pluvialis* and *C. reinhardtii*, which are already well characterized and used in food and pharmaceutical industry, is very promising regarding their use for treatment against CyHV-3.

## Declarations

### Funding

This work was funded by the German Federal Ministry of Food and Agriculture (BMEL) through the Federal Office of Agriculture and Food (BLE), grant number 2815HS010.

### Conflicts of interest/Competing interests

The authors have no relevant financial or non-financial interests to disclose.

### Availability of data and material (data transparency)

All data generated or analyzed during this study are included in this published article and its supplementary information files.

### Consent for publication (include appropriate statements)

All authors contributed to the study conception and design. Material preparation, data collection and analysis were performed by Stefanie Fritzsche. The first draft of the manuscript was written by Stefanie Fritzsche and all authors commented on previous versions of the manuscript. All authors read and approved the final manuscript.

## Acknowledgments

We would like to acknowledge Dr. Peiyu Lee and Dr. Dr. habil. Sven M. Bergmann for providing the CyHV-3 isolate that was used in this study.

---

**Table S1:** Overview of the microorganisms used for antiviral screening with information about the source and cultivation medium.

| Species | Division | Source, strain number | Cultivation conditions (medium, T, CO<sub>2</sub>, PFD) |
| --- | --- | --- | --- |
| *A. platensis* | Cyanobacteria | NIES-39 | modified *A. platensis* medium (Baer 2018), 35 °C, 3 % CO<sub>2</sub>, 180 µmol m<sup>-2</sup> s<sup>-1</sup> |
| *N. punctiforme* | Cyanobacteria | SAG 69.79 | BG11, 25 °C, 3 % CO<sub>2</sub>, 40 µmol m<sup>-2</sup> s<sup>-1</sup> |
| *C. reinhardtii* | Chlorophyta | CC-125 | TAP, 22 °C, 2.6 % CO<sub>2</sub>, 100 µmol m<sup>-2</sup> s<sup>-1</sup> |
| *C. kessleri* | Chlorophyta | SAG 211-11g | BG11, 25 °C, 3 % CO<sub>2</sub>, 40 µmol m<sup>-2</sup> s<sup>-1</sup> |
| *H. pluvialis* | Chlorophyta | SAG 34-1a | M1b, 25 °C, 3 % CO<sub>2</sub>, 40 µmol m<sup>-2</sup> s<sup>-1</sup> |
| *S. obliquus* | Chlorophyta | SAG 276-3d | M1b, 25 °C, 3 % CO<sub>2</sub>, 40 µmol m<sup>-2</sup> s<sup>-1</sup> |

CC: Chlamydomonas Resource Center, University of Minnesota NIES: Microbial Culture Collection at the National Institute for Environmental Studies, Japan SAG: Culture Collection of Algae at Göttingen University PFD: photon flux density

**Table S2:** Thermal cycling protocol used for real-time PCR for the quantitative detection of CyHV-3 copy numbers.

| Step | Temperature | Duration |
| --- | --- | --- |
| 1 Polymerase Activation and DNA Denaturation | 95 °C | 5 min |
| 2 Denaturation | 95 °C | 5 sec |
| 3 Annealing / Extension and Plate Read | 60 °C | 30 sec |
| 40 Cycles (Repetition of step 2 and 3) |  |  |

**Table S3:**
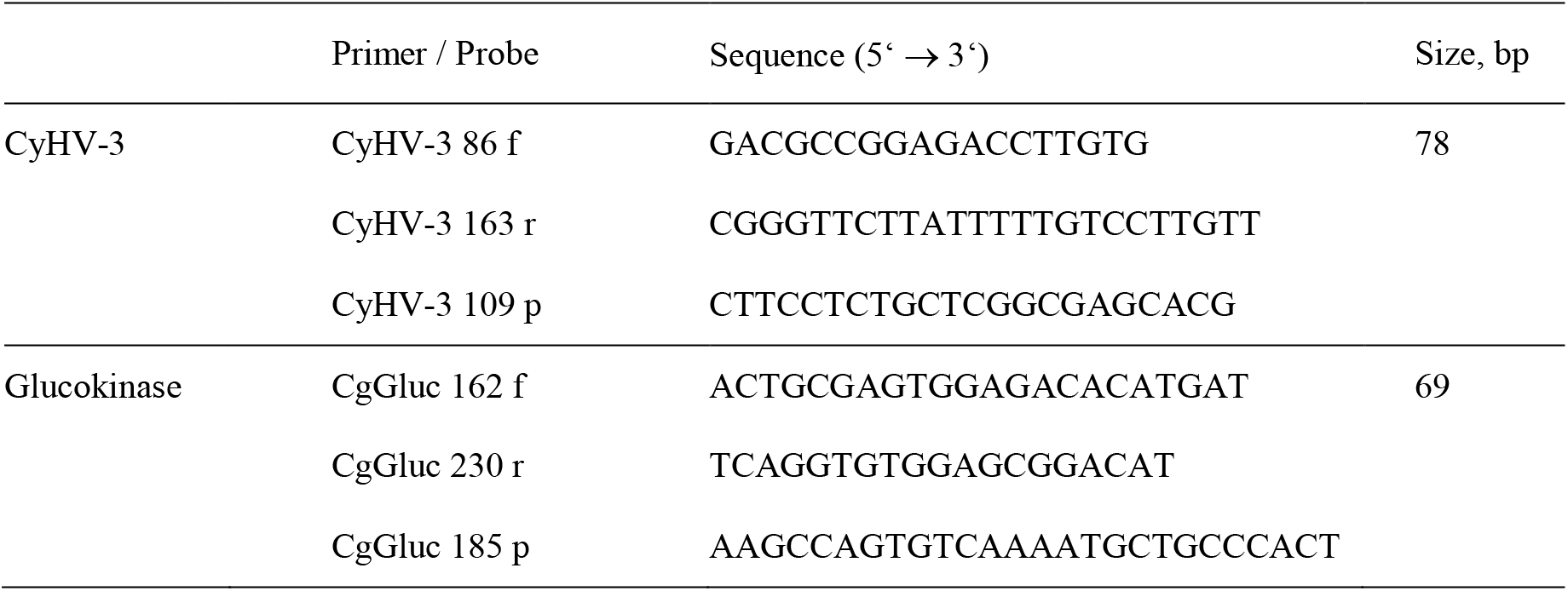
Overview of the primers and probes used for the quantitative detection of viral and cellular DNA using qPCR according to Gilad et al. (2004).

**Figure S1:**
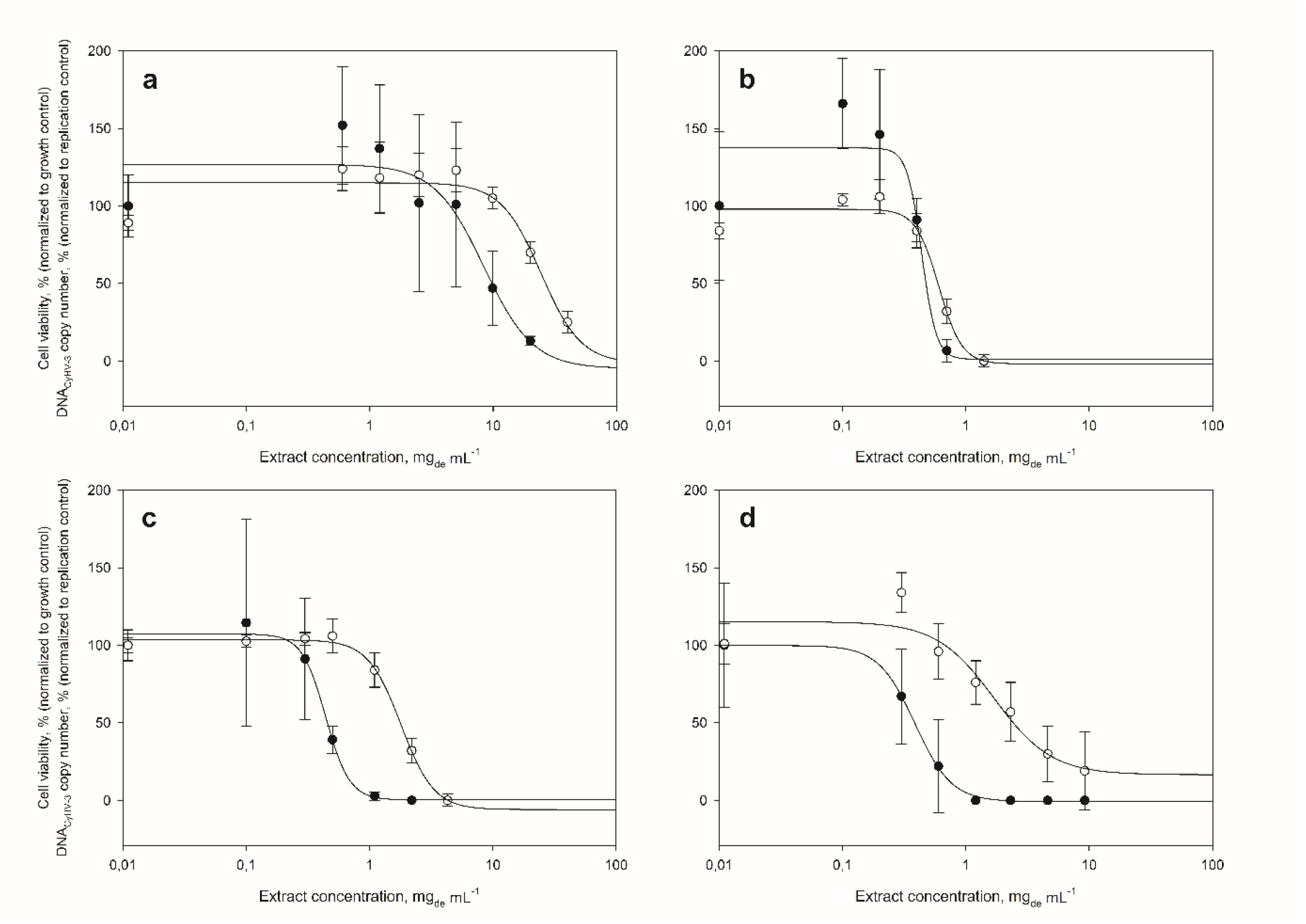
Cytotoxic and virus inhibiting effects of basic *C. kessleri* (a), *S. obliquus* (b), *N. punctiforme* (c) and *C. reinhardtii* (d) extracts, shown as normalized cell viability (white circles, based on growth control) and normalized CyHV-3 copy number (black circles, based on replication control) depending on the extract concentration. The values shown are the mean values of six biological replicates ± SD. Dose-response curves were approximated by regression using the *Four Parameter Logistic Curve* (Sigma Plot).

**Figure S2:**
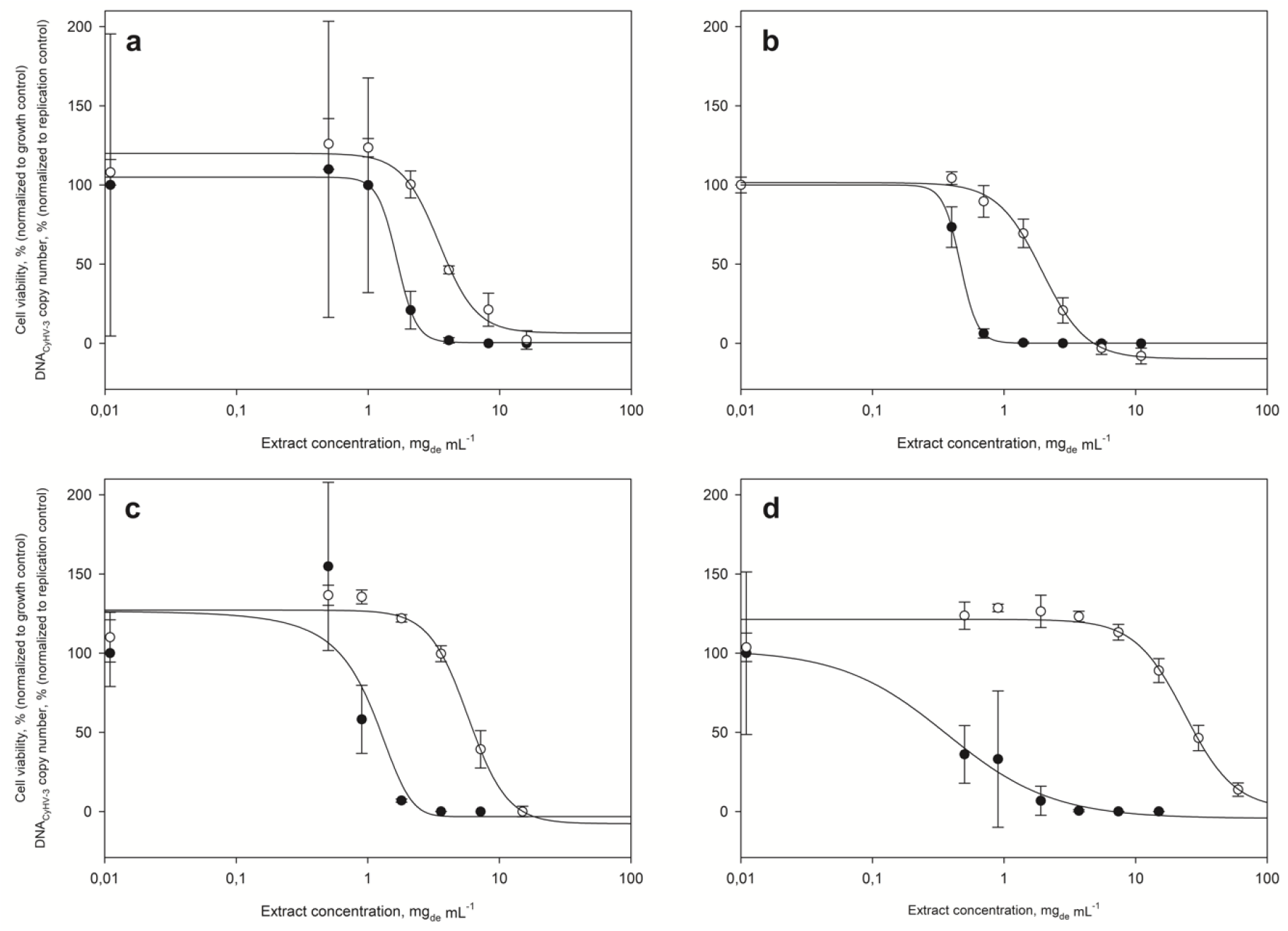
Cytotoxic and virus inhibiting effects of enhanced *A. platensis* (a), *N. punctiforme* (b), *C. reinhardtii* (c) and *H. pluvialis* (d) extracts, shown as normalized cell viability (white circles, based on growth control) and normalized CyHV-3 copy number (black circles, based on replication control) depending on the extract concentration. The values shown are the mean values of six biological replicates ± SD. Dose-response curves were approximated by regression using the *Four Parameter Logistic Curve* (Sigma Plot).

Interestingly, especially in samples containing low extract concentrations, normalized cell viabilities of above 100 % were determined using the MTT assay, suggesting viabilities better than observed in replication controls. This phenomenon has already been found when investigating the influence of cell-toxic substances by the MTT assay which is based on the metabolic activity of the cells (Weyermann et al. 2005; Isa et al. 2019). It could be explained by the hypothesis of hormesis, constituting a positive biological response to low concentrations of toxic substances (Calabrese and Baldwin 2001). Hormesis is part of many dose-response relationships and can lead to a maximal stimulation of 30 to 60 % compared to control values (Calabrese et al. 1999). Therefore, it is possible that the same extracts showing a cytotoxic effect at high concentrations, had a slightly stimulating effect on the metabolism of the CCB cells at exceptionally low concentrations, resulting in increased viability. Another possibility is the dilution of the extracts and thus the partly different concentration of cell damaging DMSO in the samples. Both, the replication control and the undiluted extract contained 1% DMSO from the redissolution process. Since the concentration of DMSO decreased with the consecutive extract dilutions, the latter could have less cell-damaging effects at low extract concentrations compared to the replication control. Despite this and the relatively high standard deviation of this biological assay, the MTT assay enabled the evaluation of cytotoxic effects of the extracts on the host cells. After the 72 h incubation periods, the cell viability of the replication control was mostly comparable to that of the growth control. Therefore, it can be assumed that the affected vitalities of the samples were actually consequences of the addition of extract.

## Notes

### Competing Interest Statement

The authors have declared no competing interest.

## References

Ahmadi A, Zorofchian Moghadamtousi S, Abubakar S, Zandi K (2015) Antiviral Potential of Algae Polysaccharides Isolated from Marine Sources: A Review. Biomed Res Int 2015:1–10. https://doi.org/10.1155/2015/825203

Amtmann A, Ahmed I, Zahner-Rimmel P, et al (2020) Virucidal effects of various agents—including protease—against koi herpesvirus and viral haemorrhagic septicaemia virus. J Fish Dis 43:185–195. https://doi.org/10.1111/jfd.13106

Baba M, Snoeck R, Pauwels R, De Clercq E (1988) Sulfated polysaccharides are potent and selective inhibitors of various enveloped viruses, including herpes simplex virus, cytomegalovirus, vesicular stomatitis virus, and human immunodeficiency virus. Antimicrob Agents Chemother 32:1742–1745. https://doi.org/10.1128/AAC.32.11.1742

Balzarini J, Schols D, Neyts J, et al (1991) Alpha-(1-3)- and alpha-(1-6)-D-mannose-specific plant lectins are markedly inhibitory to human immunodeficiency virus and cytomegalovirus infections in vitro. Antimicrob Agents Chemother 35:410–416. https://doi.org/10.1128/AAC.35.3.410

Borowitzka MA (1995) Microalgae as sources of pharmaceuticals and other biologically active compounds. J Appl Phycol 7:3–15. https://doi.org/10.1007/BF00003544

Chen F, Suttle CA (1996) Evolutionary relationships among large double-stranded DNA viruses that infect microalgae and other organisms as inferred from DNA polymerase genes. Virology 219:170–178. https://doi.org/10.1006/viro.1996.0234

Choudhary S, Save SN, Vavilala SL (2018) Unravelling the inhibitory activity of Chlamydomonas reinhardtii sulfated polysaccharides against α-Synuclein fibrillation. Sci Rep 8:1–12. https://doi.org/10.1038/s41598-018-24079-7

Defer D, Bourgougnon N, Fleury Y (2009) Screening for antibacterial and antiviral activities in three bivalve and two gastropod marine molluscs. Aquaculture 293:1–7. https://doi.org/10.1016/j.aquaculture.2009.03.047

Fabregas J, García D, Fernandez-Alonso M, et al (1999) In vitro inhibition of the replication of haemorrhagic septicaemia virus (VHSV) and African swine fever virus (ASFV) by extracts from marine microalgae. Antiviral Res 44:67–73. https://doi.org/10.1016/S0166-3542(99)00049-2

Food and Agriculture Organization of the United Nations (2017) Aquaculture production and trade trends: carp, tilapia and shrimp – FAO Project “Strengthening capacities, policies and national action plans on prudent and responsible use of antimicrobials in fisheries. Singapore

Friedrich-Löffler-Institut (2016) Koi-Herpesvirus-Infektion der Karpfen (KHV-I) – Amtliche Methodensammlung

Gallego R, Montero L, Cifuentes A, et al (2018) Green Extraction of Bioactive Compounds from Microalgae. J Anal Test 2:109–123. https://doi.org/10.1007/s41664-018-0061-9

Gey MH (2015) Instrumentelle Analytik und Bioanalytik, 3rd edn. Springer-Verlag, Berlin

Gilad O, Yun S, Adkison MA, et al (2003) Molecular comparison of isolates of an emerging fish pathogen, koi herpesvirus, and the effect of water temperature on mortality of experimentally infected koi. J Gen Virol 84:2661–2668. https://doi.org/10.1099/vir.0.19323-0

Gilad O, Yun S, Zagmutt-Vergara FJ, et al (2004) Concentrations of a Koi herpesvirus (KHV) in tissues of experimentally infected Cyprinus carpio koi as assessed by real-time TaqMan PCR. Dis Aquat Organ 60:179–187. https://doi.org/10.3354/dao060179

Gustafson KR, Cardellina JH, Fuller RW, et al (1989) AIDS-antiviral sulfolipids from cyanobacteria (Blue-Green Algae). J Natl Cancer Inst 81:1254–1258. https://doi.org/10.1093/jnci/81.16.1254

Haetrakul T, Dunbar SG, Chansue N (2018) Antiviral activities of Clinacanthus nutans (Burm.f.) Lindau extract against Cyprinid herpesvirus 3 in koi (Cyprinus carpio koi). J Fish Dis 41:581–587. https://doi.org/10.1111/jfd.12757

Haetrakul T, Tangtrongpiros J, Suthamnajpong N, Chansue N (2010) Cytotoxicity concentration of acyclovir and Clinacanthus nutans (Burm. f.) Lindau. extract to koi fin cell line. Thai J Vet Med 40:108

Harden EA, Falshaw R, Carnachan SM, et al (2009) Virucidal activity of polysaccharide extracts from four algal species against herpes simplex virus. Antiviral Res 83:282–289. https://doi.org/10.1016/j.antiviral.2009.06.007

Hayashi K, Hayashi T, Kojima I (1996) A natural sulfated polysaccharide, calcium spirulan, isolated from Spirulina platensis: In vitro and ex vivo evaluation of anti-herpes simplex virus and anti-human immunodeficiency virus activities. AIDS Res Hum Retroviruses 12:1463–1471. https://doi.org/10.1089/aid.1996.12.1463

Hayashi K, Hayashi T, Morita N, Kojima I (1993) An extract from Spirulina platensis is a selective inhibitor of herpes simplex virus type 1 penetration into HeLa cells. Phyther Res 7:76–80. https://doi.org/10.1002/ptr.2650070118

Hedrick RP, Gilad O, Yun S, et al (2000) A Herpesvirus Associated with Mass Mortality of Juvenile and Adult Koi, a Strain of Common Carp. J Aquat Anim Health 12:44–57. https://doi.org/10.1577/1548-8667(2000)012<0044:AHAWMM>2.0.CO;2

Hernandez-Corona A, Nieves I, Meckes M, et al (2002a) Antiviral activity of Spirulina maxima against herpes simplex virus type 2. Antiviral Res 56:279–285. https://doi.org/10.1016/S0166-3542(02)00132-8

Hernandez-Corona A, Nieves I, Meckes M, et al (2002b) Antiviral activity of Spirulina maxima against herpes simplex virus type 2. Antiviral Res 56:279–285. https://doi.org/10.1016/S0166-3542(02)00132-8

Herrero M, Castro-Puyana M, Mendiola JA, Ibañez E (2013) Compressed fluids (SFE, PLE and SWE) for the extraction of bioactive compounds. J Chem Inf Model 53:1689–1699

Ishaq AG, Matias-Peralta HM, Basri H (2016) Bioactive Compounds from Green Microalga Scenedesmus and its Potential Applications: A Brief Review. Pertanika J Trop Agric Sci 39:1–16

Kamble P, Cheriyamundath S, Lopus M, Sirisha VL (2018) Chemical characteristics, antioxidant and anticancer potential of sulfated polysaccharides from Chlamydomonas reinhardtii. J Appl Phycol 30:1641–1653. https://doi.org/10.1007/s10811-018-1397-2

Kanekiyo K, Lee JB, Hayashi K, et al (2005) Isolation of an antiviral polysaccharide, nostoflan, from a terrestrial cyanobacterium, Nostoc flagelliforme. J Nat Prod 68:1037–1041. https://doi.org/10.1021/np050056c

König T (2007) Gewinnung und Charakterisierung antiviraler Wirkstoffe aus aquatischen Mikroorganismen. Dissertation, Universität Erlangen-Nürnberg

Ligor M, Ratiu IA, Kiełbasa A, et al (2018) Extraction approaches used for the determination of biologically active compounds (cyclitols, polyphenols and saponins) isolated from plant material. Electrophoresis 39:1860–1874. https://doi.org/10.1002/elps.201700431

Lucero BDA, Gomes CRB, Frugulhetti ICDPP, et al (2006) Synthesis and anti-HSV-1 activity of quinolonic acyclovir analogues. Bioorganic Med Chem Lett 16:1010–1013. https://doi.org/10.1016/j.bmcl.2005.10.111

Luthria D, Luthria D, Vinjamoori D, et al (2004) Accelerated Solvent Extraction. In: Oil Extraction and Analysis, 1st edn. AOCS Publishing

Mattos BB, Romanos MT V., de Souza LM, et al (2011) Glycolipids from macroalgae: Potential biomolecules for marine biotechnology? Brazilian J Pharmacogn 21:244–247. https://doi.org/10.1590/S0102-695X2011005000056

Mletzko A, Amtmann A, Bergmann S, et al (2017) Inoculation of cyprinid herpesvirus 3 (CyHV-3) on common carp brain cells—influence of process parameters on virus yield. Vitr Cell Dev Biol – Anim 53:579–585. https://doi.org/10.1007/s11626-017-0170-1

Modrow S, Falke D, Truyen U, Schätzl H (2010) Molekulare Virologie. Spektrum Akademischer Verlag, Heidelberg

Molino A, Rimauro J, Casella P, et al (2018) Extraction of astaxanthin from microalga Haematococcus pluvialis in red phase by using generally recognized as safe solvents and accelerated extraction. J Biotechnol 283:51–61. https://doi.org/10.1016/j.jbiotec.2018.07.010

Nandasiri R, Eskin NAM, Thiyam-Höllander U (2019) Antioxidative Polyphenols of Canola Meal Extracted by High Pressure: Impact of Temperature and Solvents. J Food Sci 84:3117–3128. https://doi.org/10.1111/1750-3841.14799

Naumann I (2009) Sulfoquinovosyldiacylglyceride – antiviral aktive Substanzen. Dissertation, Universität Erlangen-Nürnberg

Neukirch M, Böttcher K, Bunnajirakul S (1999) Isolation of a virus from koi with altered gills. Bull Eur Assoc Fish Pathol 19:221–224

Norton TA, Melkonian M, Andersen RA (1996) Algal biodiversity. Phycologia 35:308–326. https://doi.org/10.2216/i0031-8884-35-4-308.1

Nuhu AA (2013) Spirulina (Arthrospira): An Important Source of Nutritional and Medicinal Compounds. J Mar Biol 2013:1–8. https://doi.org/10.1155/2013/325636

Perelberg A, Smirnov M, Hutoran M, et al (2003) Epidemiological description of a new viral disease afflicting cultured Cyprinus carpio in Israel. Isr J Aquac – Bamidgeh 55:5–12

Plaza M, Santoyo S, Jaime L, et al (2012) Comprehensive characterization of the functional activities of pressurized liquid and ultrasound-assisted extracts from Chlorella vulgaris. LWT – Food Sci Technol 46:245–253. https://doi.org/10.1016/j.lwt.2011.09.024

Pokorova D, Vesely T, Piackova V, et al (2005) Current knowledge on koi herpesvirus (KHV): A review. Vet Med (Praha) 50:139–148. https://doi.org/10.17221/5607-VETMED

Pulz O, Gross W (2004) Valuable products from biotechnology of microalgae. Appl Microbiol Biotechnol 65:635–648. https://doi.org/10.1007/s00253-004-1647-x

Rechter S, König T, Auerochs S, et al (2006) Antiviral activity of Arthrospira-derived spirulan-like substances. Antiviral Res 72:197–206. https://doi.org/10.1016/j.antiviral.2006.06.004

Reichert M (2016) Antiviral Substances from Microalgae Applied in Aquacultures. Anwendung von Antiviralen Substanzen aus Mikroalgen in Aquakulturen. Lehrstuhls für Bioverfahrenstechnik

Reichert M, Bergmann SM, Hwang J, et al (2017) Antiviral activity of exopolysaccharides from Arthrospira platensis against koi herpesvirus. J Fish Dis 40:1441–1450. https://doi.org/10.1111/jfd.12618

Ronen A, Perelberg A, Abramowitz J, et al (2003) Efficient vaccine against the virus causing a lethal disease in cultured Cyprinus carpio. Vaccine 21:4677–4684. https://doi.org/10.1016/S0264-410X(03)00523-1

Santoyo S, Jaime L, Plaza M, et al (2012) Antiviral compounds obtained from microalgae commonly used as carotenoid sources. J Appl Phycol 24:731–741. https://doi.org/10.1007/s10811-011-9692-1

Santoyo S, Plaza M, Jaime L, et al (2010) Pressurized liquid extraction as an alternative process to obtain antiviral agents from the edible microalga Chlorella vulgaris. J Agric Food Chem 58:8522–8527. https://doi.org/10.1021/jf100369h

Santoyo S, Rodríguez-Meizoso I, Cifuentes A, et al (2009) Green processes based on the extraction with pressurized fluids to obtain potent antimicrobials from Haematococcus pluvialis microalgae. LWT – Food Sci Technol 42:1213–1218. https://doi.org/10.1016/j.lwt.2009.01.012

Scott Chialvo CH, Griffin LH, Reed LK, Ciesla L (2020) Exhaustive extraction of cyclopeptides from Amanita phalloides: Guidelines for working with complex mixtures of secondary metabolites. Ecol Evol 10:4233–4240. https://doi.org/10.1002/ece3.6191

Singab AN, Ibrahim N, Elsayed AE, et al (2018) Antiviral, cytotoxic, antioxidant and anti-cholinesterase activities of polysaccharides isolated from microalgae Spirulina platensis, Scenedesmus obliquus and Dunaliella salina. Arch Pharm Sci Ain Shams Univ 2:121–137. https://doi.org/10.21608/aps.2018.18740

Thromar H, Isaacs CE, Kim K, Brown HR (1994) Inactivation of Visna Virus and Other Enveloped Viruses by Free Fatty Acids and Monoglycerides. Ann N Y Acad Sci 724:465–471. https://doi.org/10.1111/j.1749-6632.1994.tb38948.x

Thulke S (2007) Screening und Charakterisierung von Anti-beta-Herpesvirus-Aktivitäten in Substanzen aus phototrophen, aquatischen Mikroorganismen. Dissertation, Technische Universität Berlin

Tolivia A, Conforti V, Córdoba O, Flores L (2013) Chemical constituents and biological activity of Euglena gracilis extracts. J Pharm Res 7:209–214. https://doi.org/10.1016/j.jopr.2013.03.019

Troszok A, Kolek L, Szczygieł J, et al (2018) Acyclovir inhibits Cyprinid herpesvirus 3 multiplication in vitro. J Fish Dis 41:1709–1718. https://doi.org/10.1111/jfd.12880

Villarreal EC (2001) Current and potential therapies for the treatment of herpes-virus infections. Prog Drug Res 56:77–120. https://doi.org/10.1007/978-3-0348-8319-1_2

Vishwakarma J, Parmar V, Vavilala S (2019) Nitrate stress-induced bioactive sulfated polysaccharides from Chlamydomonas reinhardtii. Biomed Res J 6:7. https://doi.org/10.4103/bmrj.bmrj_8_19

Wang Y, Zeng W, Li Y, et al (2015) Development and characterization of a cell line from the snout of koi (Cyprinus carpio L.) for detection of koi herpesvirus. Aquaculture 435:310–317. https://doi.org/10.1016/j.aquaculture.2014.10.006

Wildman HG (2003) The fall and rise of natural products screening for drug discovery. Fungal Divers 13:221–231

Witvrouw M, De Clercq E (1997) Sulfated polysaccharides extracted from sea algae as potential antiviral drugs. Gen Pharmacol 29:497–511. https://doi.org/10.1016/S0306-3623(96)00563-0

Yao L, Gerde JA, Lee SL, et al (2015) Microalgae lipid characterization. J Agric Food Chem 63:1773–1787. https://doi.org/10.1021/jf5050603

## References

Baer S (2018) Screening und Optimierung von Produktionsorganismen für mikroalgenbasierte Bioraffinerieverfahren. Dissertation, Universität Erlangen-Nürnberg

Calabrese EJ, Baldwin LA (2001) Hormesis: A generalizable and unifying hypothesis. Crit Rev Toxicol 31:353–424. https://doi.org/10.1080/20014091111730

Calabrese EJ, Baldwin LA, Holland CD (1999) Hormesis: A highly generalizable and reproducible phenomenon with important implications for risk assessment. Risk Anal 19:261–281. https://doi.org/10.1023/A:1006977728215

Isa HI, Ferreira GCH, Crafford JE, Botha CJ (2019) Epoxyscillirosidine induced cytotoxicity and ultrastructural changes in a rat embryonic cardiomyocyte (H9c2) cell line. Toxins (Basel) 11:1–11. https://doi.org/10.3390/toxins11050284

Weyermann J, Lochmann D, Zimmer A (2005) A practical note on the use of cytotoxicity assays. Int J Pharm 288:369–376. https://doi.org/10.1016/j.ijpharm.2004.09.018

